# GNNQQNY: Methodology for biophysical and structural understanding of aggregation

**DOI:** 10.1101/2022.01.01.474692

**Authors:** Gunasekhar Burra, Mahmoud B. Maina, Louise C. Serpell, Ashwani K. Thakur

## Abstract

GNNQQNY sequence offers crucial information about the formation and structure of an amyloid fibril. In this study, we demonstrate a reproducible solubilisation protocol where the reduction of pH to 2.0 resulted in the generation of GNNQQNY monomers. The subsequent ultracentrifugation step removes the residual insoluble peptide from the homogeneous solution. This procedure ensures and allows the peptides to remain monomers till their aggregation is triggered by adjusting the pH to 7.2. The aggregation kinetics analysis showed a distinct lag-phase that is concentration-dependent, indicating nucleation-dependent aggregation kinetics. Nucleation kinetics analysis suggested a critical nucleus of size ∼7 monomers at physiological conditions. The formed nucleus acts as a template for further self-assembly leading to the formation of highly ordered amyloid fibrils. These findings suggest that the proposed solubilisation protocol provides the basis for understanding the kinetics and thermodynamics of amyloid nucleation and elongation in GNNQQNY sequences. This procedure can also be used for solubilising such small amyloidogenic sequences for their biophysical studies.

## Introduction

Peptide fragments are widely used for physico-chemical understanding of the otherwise complex mechanism of *in vivo* protein aggregation[1, 2]. The relatively lower complexities offered by these fragments makes them ideal models for *in vitro* aggregation studies[3-5]. The N-terminal ^7^GNNQQNY^13^ sequence of Sup35 protein serves to be useful for understanding the biophysical and structural aspects of amyloids[6]. X-ray fibre diffraction analysis of amyloid-like nano-crystals and fibrils formed by GNNQQNY showed characteristic reflections of a cross-β spine containing ‘Steric-zipper’ structure[3, 7-12]. Significant progress was achieved so far in understanding the molecular architecture of mature amyloid fibers formed by GNNQQNY, which were further complemented by simulation studies[13, 14]. But, the early steps of *in vitro* kinetic and thermodynamic changes during its self-assembly are yet to be elucidated.

Much of the information regarding the early steps of amyloid formation by GNNQQNY comes from the computer simulation studies. These studies elucidated the thermodynamic and kinetic details associated with the transition of GNNQQNY monomers to oligomers to mature fibrils[13-21]. Some studies even suggested that there exists an enthalpic barrier step, pointing towards the nucleation-dependent aggregation mechanism through which it forms fibrils rich in β-sheets[2, 10, 11, 13, 15-18, 22, 23]. This involves the formation of a metastable critical nucleus within the monomer pool along the path of aggregation. The formed nucleus acts as a template for further monomer addition at a much faster rate leading to the elongation of amyloid fibrils[24-27].

Nelson *et al* (2005) predicted that 3-4 GNNQQNY monomers constitute the metastable nucleus based on theoretical energetics[10]. Molecular dynamic simulations study by Nasica-Labouze and Mousseau (2012) suggested a nucleus of size about 4-5 monomers and 5-6 monomers at 280 K and 300 K temperature, respectively[23]. In a recent study, Szała-Mendyk and Molski (2020) suggested a relatively larger critical nucleus of size ∼25 monomers[20]. In contrast to these observations, Langkilde *et al* (2015) observed lack of intermediates during the aggregation process and thus suggested that the elongation process is likely to occur *via* monomer addition[28]. However, these studies will have practical implications only when they are proven through experimental evidences.

Although significant progress was achieved in understanding the molecular architecture of mature GNNQQNY amyloid fibers, the early steps of *in vitro* kinetics and thermodynamics of its assembly still need to be elucidated. This is probably due to the lack of a reproducible solubilisation procedure that ensures the homogenous monomeric solution to begin with. Usually the presence of insoluble aggregates due to improper solubilisation acts as templates for further monomer addition and the variation in the starting conformations leads to conflicting results[29, 30].

To overcome this issue, solvent-based pre-processing methods were usually employed to improve the solubility of proteins and amyloidogenic sequences such as amyloid-β (Aβ), polyglutamine-containing polypeptides, insulin amyloid polypeptide (IAPP), and etc[29-39]. These methods use different solvents such as trifluoroacetic acid (TFA), Dimethyl sulfoxide (DMSO), trifluoroethanol (TFE), hexafluoroisopropanol (HFIP) and ammonium hydroxide. Our recent study on polyglutamine-containing polypeptides showed that the organic solvents, TFA and HFIP aid in solubilisation by promoting the exchange of strong and intricate side chain-side chain and side chain-main chain hydrogen bonds present within the lyophilized peptides with themselves[30]. Similar observations were reported later by Jakubek *et al* (2019) as well based on UV resonance Raman spectroscopy analysis[40].

In this study, we demonstrate a solubilisation protocol that generates homogenous monomeric solution reproducibly. This enabled the study of aggregation and nucleation kinetics analyses of GNNQQNY by quantitative RP-HPLC-based sedimentation assay. The data convincingly suggested nucleation-dependent aggregation kinetics for GNNQQNY. The nucleation kinetics analysis under physiological conditions (pH 7.4 and 37 °C) resulted in a critical nucleus of size seven monomers. The mature fibrils were long unbranched structures and stained positively for ThT and Congo-red dyes. They were also found to be non-toxic and thus caused no significant cell death as compared to controls.

## Materials and Methods

### Materials

Pure form of GNNQQNY (>95%), Trifluoroacetic acid (TFA), formic acid, Congo-red and thioflavin T were obtained from Sigma Aldrich. HPLC-grade acetonitrile (ACN) and hydrochloric acid (HCl) were purchased from Merck and sodium-azide was procured from SD fine chemicals Ltd. Phosphate-buffered saline (PBS) was prepared as per Cold Spring Harbor protocol by dissolving Na_2_HPO_4_ (7.2 g), KH_2_PO_4_ (1.2 g), NaCl (40 g) and KCl (1 g) procured from Merck.

### Solubilisation of peptides

GNNQQNY peptide was solubilised at 2 mg/mL concentration in milliQ-water acidified to pH 2.0 using TFA or HCl. The solution was gently swirled till the white precipitate disappears to ensure complete solubilisation of the peptide. The resultant solution was then subjected to ultracentrifugation for a minimum of 30 min to 2 h at 80,000 RPM and 25 ^°^C. Carefully the top 2/3^rd^ of the supernatant was taken, leaving behind the bottom 1/3^rd^ of the solution which may contain the insoluble aggregates. This supernatant was used for carrying the biophysical and aggregation analysis reported in this study.

### Mass spectrometry

The *m/z* spectrum of soluble GNNQQNY peptides was recorded by injecting 5 μL of sample into time-of-flight (TOF) mass analyser equipped with electrospray ionization (ESI) source (Waters Q-TOF premier HAB213). The data was plotted at a mass range of 50–3500 Da using OriginPro 8.5 data analysis and graphing software. The purity and solubility was determined by analysing all the observed molecular ionization species of the peptide[41].

### Concentration determination by Reversed-Phase HPLC

GNNQQNY peptide stock solution (0.4 mg/mL) was prepared by solubilising as previously mentioned and used for determining the standard curve. The stock was diluted serially and the optical density (OD) of each dilution was determined at 220 nm by using NanoDrop ND-1000 UV-Vis Spectrophotometer. The corresponding concentration (μg) was back calculated based on the standard procedure suggested by Kuipers and Gruppen (2007)[29, 30, 42, 43]. Briefly, the molar extinction coefficient used for determining the concentration was obtained by summing up the molar extinction coefficients of all the amino acids and the peptide bonds at 214 nm[42]. Each of these dilutions was subjected to RP-HPLC analysis by passing through Agilent eclipse plus C_18_ column (4.6 mm × 100mm) connected to Agilent 1260 Infinity Quaternary system. A gradient flow of water and acetonitrile containing 0.05% (v/v) TFA at a rate of 1 mL/min was used for elution. The standard-curve was obtained by plotting the area under the curve determined at 214 nm for each of the dilution against the respective concentration (μg) calculated based on OD. The data was subjected to linear curve-fit using OriginPro 8.5 graphing tool and the unknown concentration (μg) of the peptide was determined by fitting the area in the standard curve.

### Analytical size–exclusion chromatography (SEC)

The pH of 306 μM of soluble GNNQQNY peptide in water-HCl (pH 2.0) was adjusted to pH 7.4 by using PBS. 100 μL of this solution was analysed using SEC method. For this, the sample was passed through Superdex peptide 10/300GL column (GE healthcare); connected to Biologic Duoflow model Bio-Rad fast protein liquid chromatography (FPLC) system. PBS (pH 7.4) was run at a flow rate of 0.5mL/min to monitor the elution profile of the peptide. The absorbance was recorded at 215 nm while maintaining a constant pressure of ∼80 psi[30, 41].

### Aggregation kinetics analysis

The aggregation kinetics reactions were monitored in PBS at pH 7.4 and 37 °C. Sodium-azide (0.05%) was added to the reaction mixture to prevent microbial contamination[29, 34, 44, 45]. The rate of aggregation was monitored by measuring the monomer concentration as a function of time by using sedimentation assay. For this, an aliquot of sample was taken from the ongoing aggregation reaction at regular time intervals and subjected to ultracentrifugation at 25,000 rcf and 25 °C for 30 min. The supernatant was taken and adjusted to 20% formic acid before injecting into RP-HPLC. The monomer concentration was determined based on the above standard curve. Addition of formic acid was shown to slowdown aggregation as it reduces the pH of the sample significantly[34].

The sedimentation assay data was complimented by monitoring the rate of formation of higher order structures by using light scattering (LS) and Thioflavin T (ThT) assays[46, 47]. These assays were performed by taking 120 μL of ongoing aggregation reaction sample at different time points in a Quartz SUPRASIL Ultra-micro cell[30]. The emission and excitation wavelengths were set at 450 nm, and slit widths at 2.5 nm for determining the intensity of scattered light. To this, 2.5 mM ThT stock was added to a final concentration of 100 μM and the excitation and emission wavelengths were reset to 450 nm (slit width, 5 nm) and 489 nm (slit width, 5 nm), respectively to measure the ThT intensity. These assays were performed using LS 55 spectrofluorimeter (Perkin Elmer). Three consecutive spectra were averaged after blank correction for obtaining the final LS and ThT spectra.

### Seeding assay

Seeding reactions were monitored in PBS (pH 7.4) at 37 °C by adding the preformed seeds (2%, wt/wt) grown at similar conditions as described previously[29, 48]. The control reactions used for comparison were monitored without addition of preformed seeds. The drop in the monomer concentration with time was determined by using sedimentation assay. The aggregation kinetics of both the conditions was compared by plotting the monomer percentages against time (h) by using OriginPro 8.5 software.

### Nucleation Kinetics analyses

Different concentrations of freshly solubilised GNNQQNY were incubated in PBS at pH 7.4 and 37 °C. The drop in monomer concentration with time was measured at different time intervals using sedimentation assay. Monomer concentration (μM) versus time^2^ (s^2^) graphs were plotted for each concentration by collecting as many data points as possible within 20% drop in monomer concentration. Using the slopes of these time^2^ graphs, log[initial concentration (M)] versus log[slope] plots were constructed. This plot was subjected to linear curve-fit and the obtained slope was fit in the equation, slope = n* + 2, where n* indicates the size of the critical nucleus[24, 29, 34, 48, 49].

### Thioflavin T (ThT) and Congo-red staining of fibrils

Standard solutions of ThT (0.8 mg/mL) and Congo-red (1 mg/mL) prepared in MilliQ-water and filtered through 0.2 μm filter were used to stain the GNNQQNY fibers. 10 μL of peptide solution was evenly spread on a neat glass slide and allowed to air-dry. A drop of filtered ThT and Congo-red solution was added on the dry peptide film and incubated for 30 min. Excess dye was removed by washing with MilliQ-water followed by air drying for 30 min. Bright field images and the associated ThT fluorescence and Congo-red birefringence images were recorded using appropriate light filters on Leica DM2500 model fluorescent microscope[3, 50].

### Fourier-transform infrared (FTIR) spectroscopy

Soluble form of GNNQQNY and the fibril suspension were analysed using Bio-ATR II cell mounted on Bruker Tensor-27 FTIR instrument[29, 30, 51]. 300 μL of fully grown GNNQQNY fibers were washed thrice by using milliQ-water and then concentrated to 100 μL. 50 μL of sample was loaded on to the Bio-ATR II cell for analysis. The solid-state FTIR analysis was performed by loading the KBr-mixed GNNQQNY powder on to MIR-ATR cell. The final FTIR spectrum was an average of 120 scans at a resolution of 4 cm^-1^. The spectrum was auto-corrected for water vapour and the blank. The second derivative of primary spectra was reported after nine-point smoothing by Opus 7.2 spectroscopy software (Bruker Optics).

### Transmission electron microscopy (TEM)

Samples were prepared by inverting the carbon-coated side of size 300 mesh copper TEM grid on 5 μL of aggregate sample for thirty seconds. The excess sample was carefully blotted by using KimWipes and was rinsed with 10 μL of milliQ-water followed by inverting on 5 μL of 2% uranyl acetate for 5-10 seconds[29, 30, 52]. The stained grids were allowed overnight to air-dry and the electron micrographs of fibrils were recorded using FEI-Tecnai G1 12 Twin (120 KV) transmission electron microscope.

### X-ray fiber diffraction

200 μL of GNNQQNY peptide (10 mg/mL) was incubated at 37 °C for several days to generate fibers. Approximately 20 μL of this sample was placed and allowed to air-dry overnight between two wax-filled capillaries. The generated fiber was placed on a goniometer head mounted on a Rigaku X-ray diffractometer (Sevenoaks, UK) equipped with rotating anode (CuKα) and Raxis IV++ detector. The specimen-to-film distance of 50 mm or 100 mm was set for recording the data[53].

### Cytotoxicity and Cell viability assays

Undifferentiated SHSY5Y cells were differentiated using established protocol[54, 55]. The differentiated cells were incubated with GNNQQNY fibers (5 μM and 20 μM) for 48 hours. Phosphate-buffered saline (1X) and H_2_O_2_ (1 mM) were used as negative and positive controls, respectively. Towards the end of the incubation, the cells were treated with 5 μM CellROX Green Reagent (C10444, Life technologies UK) for 45 min to evaluate the level of oxidative stress. This was followed by incubation with ReadyProbes reagent (Life Technologies) for 15 min to evaluate cell death. The cells were imaged at 37 ^°^C and 5% CO_2_ on Operetta CLS high-content imaging system (PerkinElmer) by using DAPI, FITC and TRITC filters. To ensure the reproducibility of the findings, three independent experiments each containing a minimum of 5000 cells were analysed using Harmony software automated analysis algorithm provided within the system[56].

## Results and discussion

The amyloidogenic sequences such as GNNQQNY poses a challenge to unambiguously resolve the kinetic and thermodynamic properties associated with its aggregation. This is mainly due to improper or partial solubilisation leading to contamination of the aggregation reaction with pre-aggregated peptide. Existence of such forms in the starting reaction results in spontaneous aggregation which is often non-reproducible owing to the heterogeneous nature of the starting material. Probability of this phenomenon was explored by analyzing the infrared spectra of GNNQQNY powder and its solubilised form (figure 1A). Solid-state infrared spectra of lyophilized GNNQQNY powder showed peaks matching to parallel (1634 cm^-1^ and 1682 cm^-1^) and anti-parallel (1623 cm^-1^ and 1689 cm^-1^) β-sheets and random coil (1648 cm^-1^) conformations (figure 1A)[30, 57, 58]. The side chain NH_2_ bending (1610 cm^-1^) and C=O stretching (1656 cm^-1^ and 1669 cm^-1^) vibrations of Gln/Asn residues were also identified (figure 1A)[59]. The solid form of GNNQQNY also showed two more prominent peaks at 1698 cm^-1^ and 1706 cm^-1^, indicating the presence of weaker H-bond interactions in the lyophilised forms (figure 1A)[30, 59]. Overall, this data suggests that the lyophilised form of GNNQQNY is an amorphous aggregate that may or may not be soluble in aqueous solution[30].

**Figure 1:**
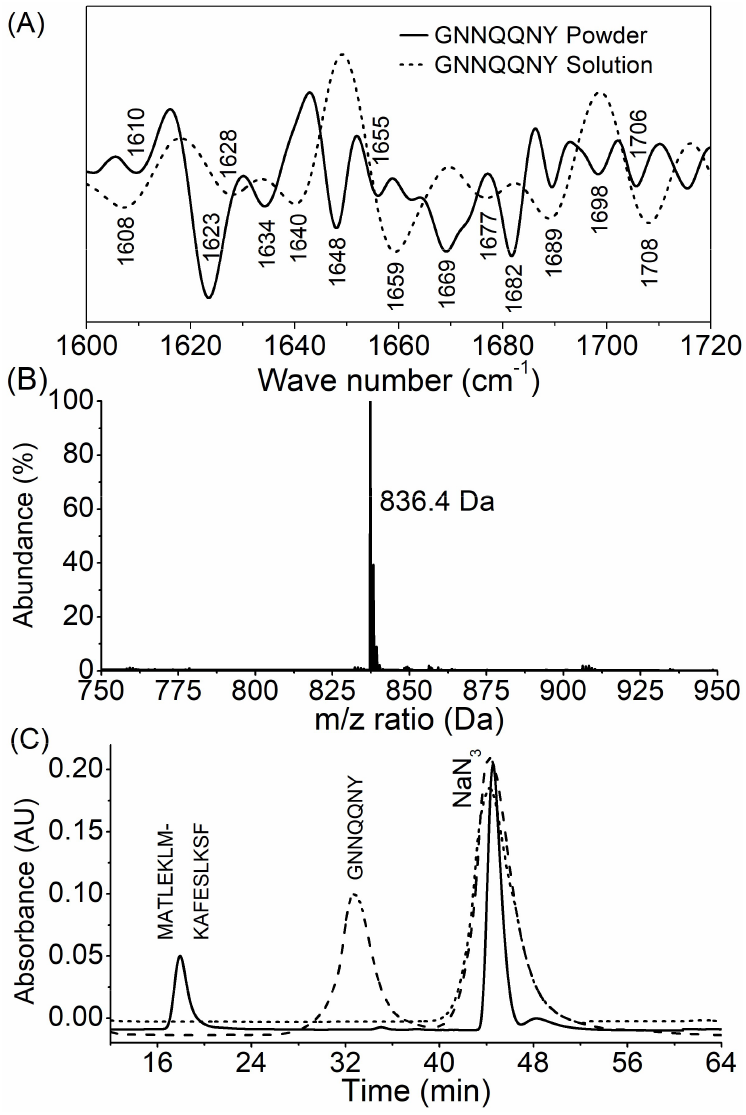
Biophysical characterization of GNNQQNY sequences. (A) Infrared spectroscopy analysis of GNNQQNY powder (solid line) indicating the amorphous conformation and GNNQQNY solution indicating dominant random coil conformation. (B) ESI-MS analysis showing the soluble monomeric (836.4 Da) state of GNNQQNY. (C) Size-exclusion chromatography analysis of GNNQQNY monomer (dash line), MATLEKLMKAFESLKSF (solid line) and sodium azide (NaN_3_, dotted line) confirming the monomeric state of the soluble fraction.

Unlike other amyloidogenic sequences such as Aβ, polyglutamines and IAPP that require special solvent pre-treatment to enhance solubility, GNNQQNY was freely soluble in water and the resultant solution was clear and transparent. When this solution was analysed using Bio-ATR IR, the characteristic bands indicating the presence of β-sheets(1623 cm^-1^, 1634 cm^-1^, 1682 cm^-1^ and 1689 cm^-1^)in the lyophilised forms were greatly diminished (figure 1A)[30, 57, 59]. In addition, the emergence of new bands suggesting a more unordered conformation (1628 cm^-1^, 1640 cm^-1^ and 1677 cm^-1^) were observed (figure 1A)[30, 57, 59]. The structural transition of β-sheets in solid form to unordered conformation in solution further confirms solubility of GNNQQNY sequences. The probable reasons for such easy transition of conformations might be the dominance of relatively less stable parallel β-sheets due to longer and less optimal H-bonds and the existence of weaker H-bond interactions that are easily accessible to the solvent[60, 61].

However, the major issue is that the GNNQQNY sequences solubilised in pure water tends to spontaneously aggregate, and thus result in non-homogeneous solution by the time the aggregation reactions are initiated. This is probably due to the fact that the isoelectric point of GNNQQNY (pI, 5.9) is close to that of the water (pI, 6.5-7.0), and that might promote its spontaneous aggregation or precipitation. One way to overcome this phenomenon is by maintaining the pH of the solution at a point that is far away from the pI of GNNQQNY. This can be achieved by either significantly increasing or decreasing the pH from 5.9. Usually, the peptides/proteins are positively charged at a pH below their pI and are negatively charged at a pH above their pI depending on their sequence[60, 61]. This property was taken advantage here and the pH of the solvent (milliQ water) was significantly reduced to 2.0 by using HCl or TFA. This point was considered owing to the fact that five out of the seven residues in GNNQQNY are polar in nature. At such low pH (2.0), the side-chains of asparagine (N) and glutamine (Q) residues are highly positively charged and thus repel with each other, thereby aiding in their solubility through interaction with water[43, 62].

Further, the homogeneous nature of the monomeric solution was ensured by subjecting the solubilised solution to ultracentrifugation (80,000 RPM) for at least 30 min at 25 °C. We have shown recently in one of our studies that temperature plays a prominent role in regulating the aggregation of GNNQQNY[43]. Particularly, the efficiency of nucleation was found to be very high at low temperatures[43]. Hence the ultracentrifugation at low temperatures and the storage of solubilised peptide under freezing conditions should be avoided for better reproducibility of results. The purity, identity and monomeric state of GNNQQNY were confirmed by native ESI-MS analysis where only the expected *m/z* value of monomer (836.4 Da) was observed (figure 1B). Subjecting the supernatant to analytical SEC analysis also resulted in a single peak, thus confirming the homogenous nature of the solution (figure 1C). Absence of additional peaks and the elution of GNNQQNY (836.4 Da) before sodium azide (NaN_3_, 65 Da) and after MATLEKLMKAFESLKSF (1974.4 Da) further confirmed the monomeric state of the solubilised fraction (figure 1C).

The lack of an aromatic amino acid residue tryptophan in GNNQQNY sequence makes it difficult to monitor the aggregation kinetics based on quantitative methods like analytical reversed-phase high performance liquid chromatography (RP-HPLC). The data generated by Thioflavin T (ThT) and light scattering assays has no practical meaning as it is qualitative in nature. Hence we took the work of Kuirper and Gruppen as basis and developed a quantitative method to follow the aggregation kinetics. In this method, the overall contribution of the molar extinction coefficients of individual amino acids and the peptide bond at 215 nm was taken into consideration for determining the concentration[42]. Upon plotting the concentration against the respective area under the curve obtained from RP-HPLC for different dilutions at 215 nm, we obtained a standard curve (figure 2A)[42]. This standard curve was used for determining the drop in monomer concentration during aggregation kinetics analysis.

**Figure 2:**
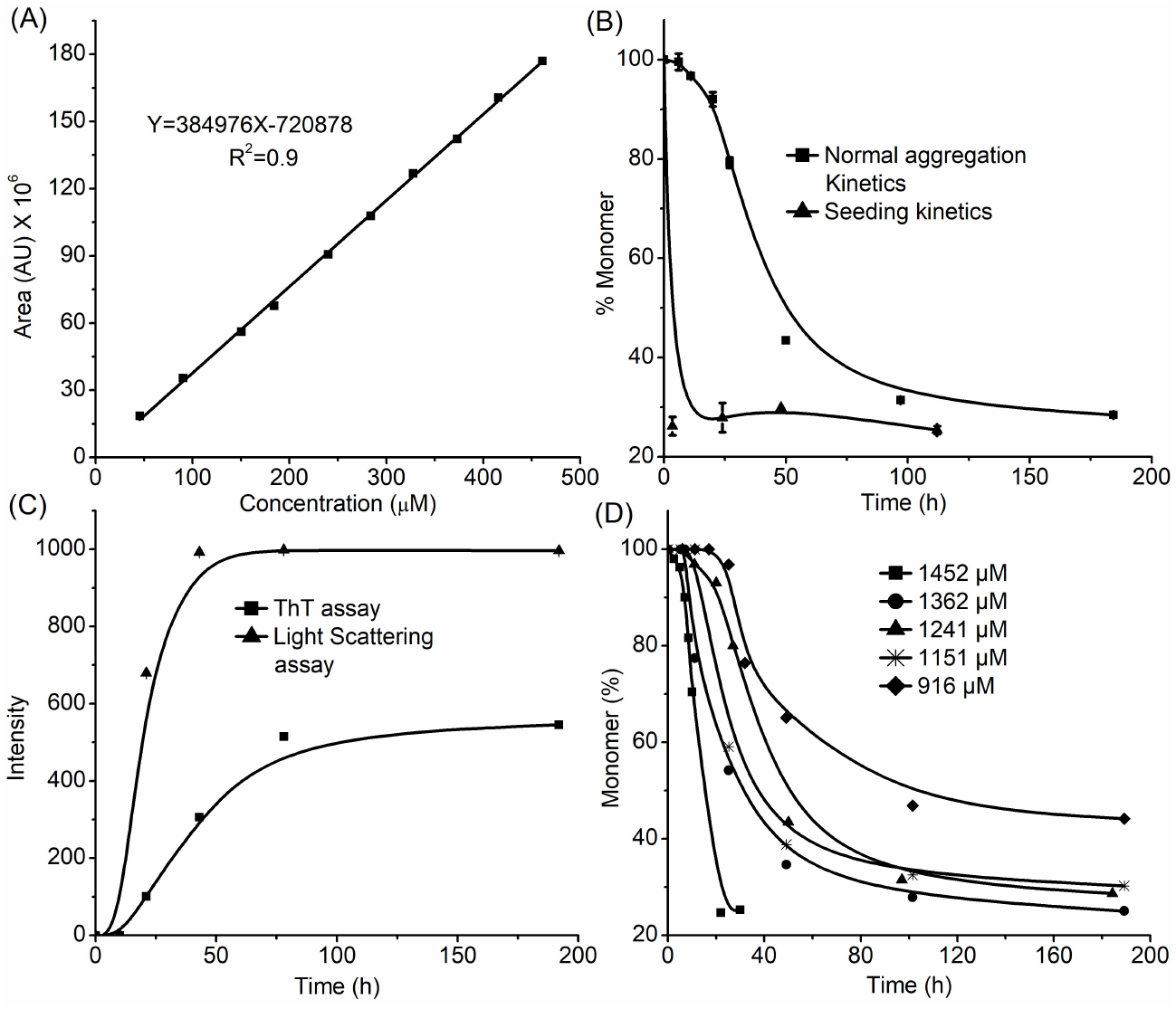
Aggregation analysis of GNNQQNY (1253±12 μM) at pH 7.4 and 37 °C. (A) Standard curve for quantifying the monomer concentration based on sedimentation assay. (B) Aggregation kinetics analyses of normal unseeded (■) and seeded (▲) reactions monitored by sedimentation assay. (C) Aggregation kinetics monitored by ThT (▲) and light scattering (■) assays showing nucleation-dependent aggregation. (D) Concentration-dependent aggregation kinetics analysis. Error bars specify the standard deviation calculated from three independent reactions.

We then tested the ability of this solubilisation method to understand the aggregation kinetics and reproduce the data reported in the literature by adjusting the pH of the sample to 7.4 at 37 °C. The rate of aggregation was monitored by measuring the drop in monomer concentration as a function of time by using sedimentation assay. This data was further supplemented by measuring the rate of formation of aggregates with time by using ThT and light scattering assays. The sedimentation assay data clearly showed a lag-phase of ∼12 h, where the drop in monomer concentration was negligible and took about 25 h for an initial drop in 20% monomer (figure 2B). Collectively this data indicated a lag-phase of 20-25 h for GNNQQNY (1253±12 μM), within which the energetically unfavourable nuclei are formed. This was followed by a rapid decrease in the concentration of monomer, indicating an elongation-phase. No further drop in monomer concentration was observed towards the end of elongation-phase, and is suggestive of an equilibrium or stationary-phase (figure 2B). The concentration of monomer at the equilibrium-phase is usually termed as the critical concentration (C_r_), below which the efficiency of nucleation is negligible. Overall, this data hinted at a nucleation-dependent aggregation mechanism for GNNQQNY[34, 49, 63, 64].

However, a classical nucleation-dependent aggregation kinetics should also display; seeding effect, specific ThT fluorescence indicating critical nucleus formation, and concentration-dependence of rate of aggregation[34, 65]. The effect of seeding on aggregation was tested by adding the preformed seeds (2%, wt/wt) at the start of the reaction. Addition of seeds eliminated the lag-phase by spontaneously reducing the monomer concentration to 25% within 3-4 h of incubation (figure 2B). This is mainly because the added aggregates act as templates for monomer addition, thereby drives the aggregation via the elongation-phase[29, 34, 63-67].

In agreement with the sedimentation assay data, an enhancement in ThT fluorescence was observed only after 12 h of incubation. The binding of ThT to aggregates usually results in enhanced fluorescence at 489 nm[68-70]. This data confirms the lack of formation of ThT positive structures during the lag-phase (figure 2C). This was followed by fluorescence enhancement along the elongation-phase and it reached the equilibrium at the stationary-phase. Observation of a similar trend by LS assay that indicates the formation of higher-order structures with time further complements the conclusions drawn from sedimentation and ThT data (figure 2B and C).

The dependence of rate of aggregation on reaction concentration was established by determining the aggregation kinetics at different starting concentrations. This data showed an inverse relation between the starting concentration of the reaction and the corresponding lag-phase observed for each concentration (figure 2D). Similarly, the critical concentration observed at the equilibrium-phase was increasing upon reducing the starting concentration of the aggregation reaction (figure 2D). These observations confirm the fact that the efficiency of nucleation in a nucleation-dependent aggregation decreases upon reducing the starting concentration[2, 24, 34, 49, 64, 65].

The phenomenon of nucleation was further confirmed by determining the actual size of the critical nucleus formed during aggregation. For this, the aggregation kinetics at different initial concentrations was monitored (figure 3A). The initial 20% drop in the monomer concentration was used to determine the concentration (μM) versus time^2^ (Sec^2^) curves (figure 3B). Plotting the log[slope] of each linear time^2^ plot against the respective log[concentration (M)] resulted in a linear log-log plot with a slope of 9.2 (figure 3C). This suggested a critical nucleus of size seven (n^*^=7, after round off) monomers. Collectively, these observations provide strong evidence for a nucleation-dependent aggregation mechanism for GNNQQNY[6, 10, 11, 22, 23].

**Figure 3:**
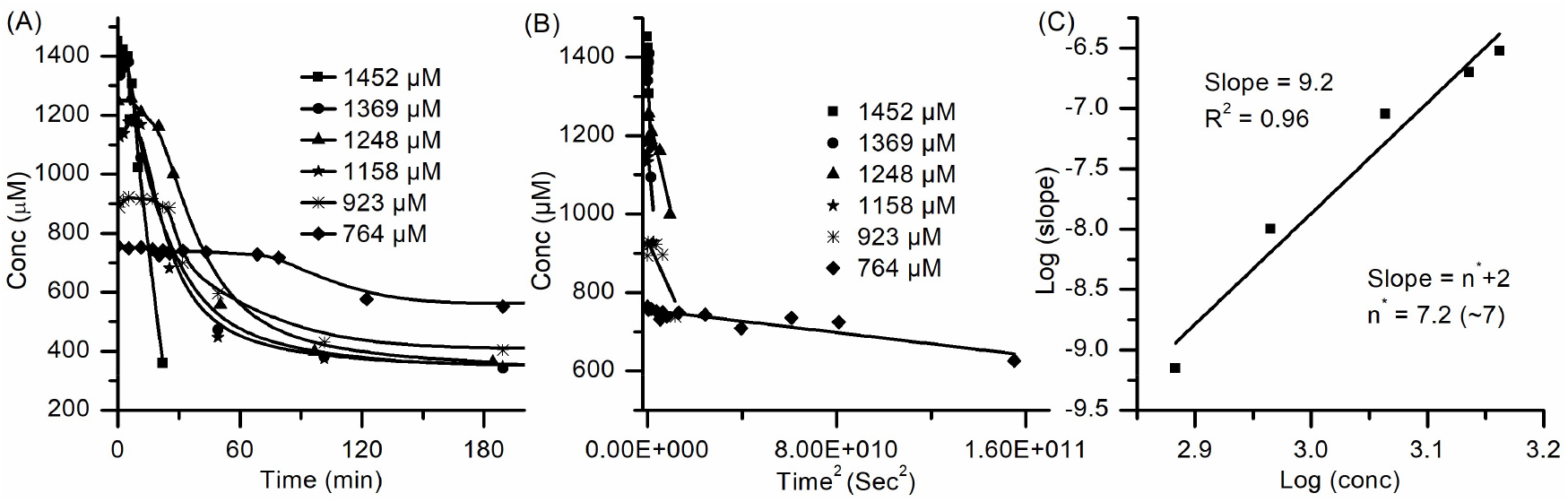
Nucleation kinetics of GNNQQNY at pH 7.4 and 37 °C. (A) Shows the Concentration-dependent aggregation kinetics of GNNQQNY at six different starting concentrations. (B) Plot of the initial 20% drop in the monomer against time^2^ (s^2^) leading to linear time^2^ plot. (C) The log-log plot obtained by plotting the log[slope] obtained from (B) against the respective log[initial concentration (M)].

The amyloid-like characteristics of mature fibrils formed at the end of aggregation were established by using different biophysical and structural characterisation methods such as FTIR, ThT, Congo-red (CR), TEM and fibre-XRD analyses (figure 4). The presence of a characteristic β-sheet band at 1631 cm^-1^ and a corresponding minor band at 1690 cm^-1^ in infrared spectroscopy data indicated the presence of β-sheet-rich structures in fibers (figure 4A). The aggregates generated through seeding of the reaction also showed similar characteristics when analysed by FTIR spectroscopy (figure 4A). These fibrils exhibited *yellow-green* fluorescence on staining with ThT and *apple-green* birefringence on staining with Congo-red dyes (figure 4B, C)[71]. The ThT fluorescence emission and CR birefringence property confirmed the presence of β-sheet-rich structures and amyloidogenic nature of fibers[50, 68, 69, 72-74]. Electron microscopy images showed long fibrous-to-crystalline structures comparable to those reported earlier for GNNQQNY (figure 4D)[28, 53, 71, 75]. Fiber diffraction analysis indicated distinct diffraction signals at 4.8 Å (meridional) and 9.3 Å (equatorial), which are characteristic signals of cross-β architecture of amyloid fibrils (figure 4E)[3, 10, 11, 53, 71, 75].

**Figure 4:**
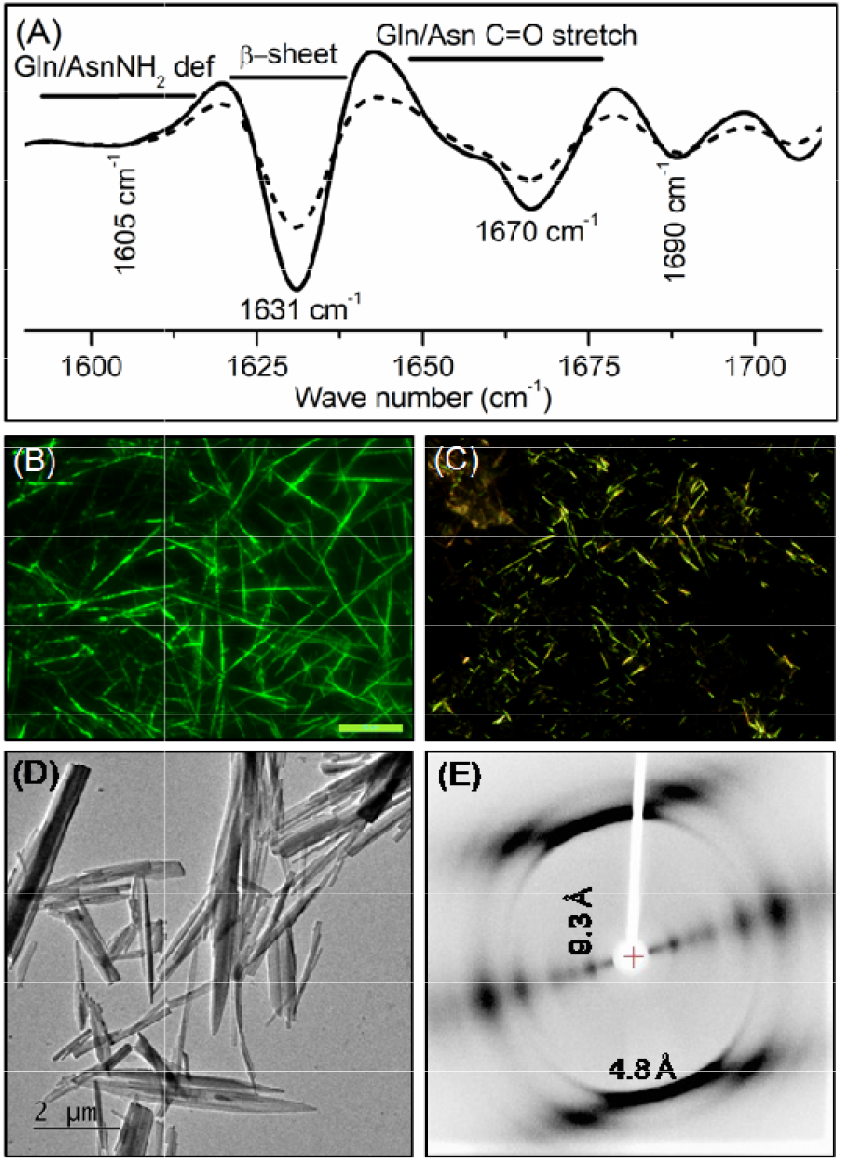
Biophysical and structural characterisation of GNNQQNY fibers. (A) FTIR spectra showing the characteristic β-sheet bands at 1631 cm^-1^ and 1691 cm^-1^. (B) ThT data showing distinct bright *yellow-green* fluorescence. (C) Congo-red staining indicating characteristic *apple-green* birefringence of amyloid-like structures. (D) Electron micrographs showing the fibrous-to-crystalline amyloid fibrils. (E) Fiber-XRD data showing the characteristic reflections fibers. confirming the cross-β spine containing amyloid cores of fibers.

GNNQQNY sequence was shown to induce the fibril formation in yeast *Sup*35 protein by constituting the fibril-core of prion fibrils[3, 76]. These fibrils being functional in nature are usually non-toxic to the yeast cells. This was confirmed by investigating the toxicity of GNNQQNY fibers by CellRox green reagent-based oxidative stress and ReadyProbes Live/Dead cell assays. Co-incubating the differentiated SHSY5Y cells with GNNQQNY aggregates at physiological conditions (pH 7.4 and 37 °C) resulted in no significant oxidative stress levels and percentage of dead cells as compared to PBS-treated cells (figure 5A, B). This outcome was also compared with that of the known inducer of oxidative stress and cell death, H_2_O_2_ for confirmation[77, 78]. As shown earlier, the cytotoxicity and cell death exhibited by H_2_O_2_ was significantly higher as compared to GNNQQNY and PBS (figure 5A, B)[30]. Altogether, these data confirmed that the fibers formed by GNNQQNY are non-toxic in nature as reported earlier[30]. All of these evidences confirmed that the aggregates generated through the proposed solubilisation method resulted in amyloid fibers that share a great similarity with the existing published literature. These striking similarities suggested the wide applicability of this method for carrying biophysical and structural studies of amyloidogenic polypeptides.

**Figure 5:**
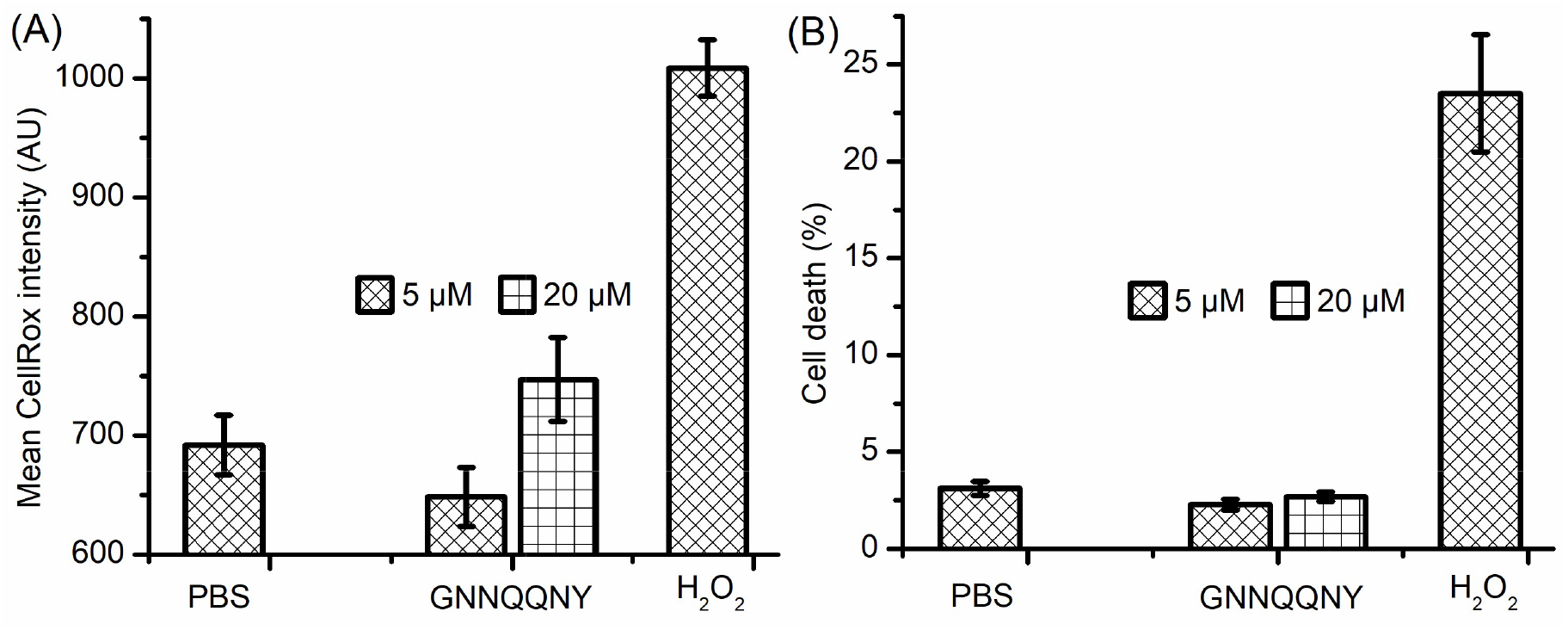
Toxicity of GNNQQNY fibers and controls. (A) Oxidative stress exhibited by GNNQQNY fibers, PBS and H_2_O_2_ on co-incubating with SHSY5Y cells for 48 h. (B) Percent cell death observed under the same conditions. Error bars indicate the standard deviation calculated based on three independent experiments. The statistical significance (*P<0.05) was determined by one-way ANOVA test.

## Conclusion

In this study, we report a solubilisation protocol that generates homogenous monomeric solution. This protocol allows the monomers to remain stable until the aggregation reaction is triggered, thus facilitating the aggregation and nucleation kinetics analyses based on quantitative RP-HPLC-based sedimentation assay. The experimental results revealed that the hepta-peptide; GNNQQNY follows nucleation-dependent aggregation kinetics. The data under physiological conditions (pH 7.4 and 37 °C) suggested that ∼7 monomers constitute to form a critical nucleus that acts as a template for further monomer addition to drive the aggregation reaction forward. The mature GNNQQNY fibers formed are long, unbranched structures and displayed *apple-green* birefringence and *green-yellow* fluorescence on staining with Congo-red and thioflavin T dyes, respectively. Also, these fibrils did not exhibit any increase in cytotoxicity or cell death upon co-incubating with SHSY5Y neuroblastoma cells. This study provides the basis for experimental understanding for the previous predictions which are based on theoretical and molecular dynamic simulations. Thus we believe that this solubilisation protocol will have wider practical implication to understand the kinetics and thermodynamics of aggregation of amyloidogenic sequences.

## Author contributions

GB conceptualised the study, executed the experiments and interpreted the results. MBM contributed in executing the cytotoxicity and cell death studies. XRD data was recorded and analysed by GB and LCS. Manuscript was prepared by GB and edited by LCS and AKT. LCS and AKT supervised the project.

## Acknowledgements

GB acknowledges the European Molecular Biology Organisation (EMBO) for providing a Short-Term Fellowship award (EMBO-STF 7674) to visit Prof. Louise C. Serpell’s lab at University of Sussex, United Kingdom. LCS acknowledges funding from Alzheimer’s Society in support of MBM (AS-PG-16b-010). LCS and MBM are members of the Alzheimer’s Research UK South Coast Network and are grateful for their support. LCS is supported by BBSRC (BB/S003312/1). AKT acknowledges Department of Biotechnology (DBT/BSBE/20120020) and Indian Council of Medical Research (ICMR/BSBE/2016489), Govt. of India for the financial support.

## Abbreviations

CR: Congo red
ESI: Electrospray ionization
FPLC: Fast protein liquid chromatography
FTIR: Fourier-transform infrared spectroscopy
HCl: Hydrogen chloride
LS: Light scattering
PBS: Phosphate-buffered saline
RP-HPLC: Reversed-phase high performance liquid chromatography
SEC: Size exclusion chromatography
TEM: Transmission electron microscopy
ThT: Thioflavin T
ToF: Time-of-flight

